# An improved haplotype resolved genome reveals more rice genes

**DOI:** 10.1101/2023.12.07.570500

**Authors:** Muhammad Abdullah, Agnelo Furtado, Ardashir Kharabian Masouleh, Pauline Okemo, Robert Henry

## Abstract

The rice reference genome (Oryza sativa ssp. japonica cv. Nipponbare) has been an important resource in plant science. We now report an improved and haplotype resolved genome sequence based upon more accurate sequencing technology. This improved assembly includes regions missing in earlier genomes sequences and the annotation of more than 3,000 new genes due to greater sequence accuracy.

## Introduction

Nipponbare is a japonica rice cultivar that has been widely used as the standard reference genotype for rice ^1^. The rice (Nipponbare) genome was one of the first crop genomes to be sequenced more than 20 years ago ^2^. The 1st sequence of the rice genome was completed in 2002 and was a major milestone in the field of plant genomics by the International Rice Genome Sequencing Project, 2005 ^3^. These international collaborative efforts provided the first genome of a crop plant. The Nipponbare genome assembly contained gaps, primarily due to repetitive DNA sequences. In 2005, these gaps were estimated to be approximately 18.1 Mb in total, with the majority originating from centromeres and telomere regions. Sequencing technological advancements and ongoing research efforts, have improved the rice genome sequence over time ^4,5^. Thorough, efforts were made to improve the quality of the Nipponbare reference genome assembly in 2013 resulting in greatly enhanced accuracy of cDNA sequences and gene annotation, while it remained incomplete ^5^. In the human genome, recent significant strides have been made in assembling and characterization the previously unexplored 8% of the human genome, especially including telomere sequences ^6^.

## Results

Reference genome assemblies often contain gaps, especially regions with repetitive sequences, termed the “dark side” of the genome ^7,8^. New sequence technology allows improved assembly quality, with less gaps, leading to a more complete and accurate representation of the genome. The achievement of a higher quality and more complete reference genome will provide new insights into genomics and breeding, supporting pan-genome studies and genome wide association studies ^9^. Recently many other *Oryza* genomes have been sequenced and assembled, including indica and wild rice species ^9-11^ Most recently the Nipponbare genome sequence gaps and telomere sequence were addressed ^12,13^. Despite these advancements, a fully haplotype resolved assembly has not been reported. In this study, we have used PacBio HiFi reads to produce a more accurate genome sequence assembly. The novel genome assembly is not only almost 11.3 megabases (Mb) longer than the IRGSP-1.0 reference but also exhibits improvements in all chromosomes, including the addition of telomeric regions in all chromosomes (T2T), with the addition of fully resolved haplotypes (haplotype 1 and haplotype2, telomere-to-telomere-T2T) (Supplementary Table S1-10). Comparative analysis of annotations of the new genome (UQ_Nipponbare) and the IRGSP-1.0 reference revealed the presence of 3,050 additional genes, for which more than 95% had supporting transcript evidence (Supplementary Figure S1). These finding underscore the potential of new sequencing technologies to significantly augment reference genomes, potentially leading to more comprehensive genetic information. These results also suggest that applying advanced sequencing technologies to other established genomes may yield similar benefits, potentially enhancing our knowledge of these species. This study highlights the continuous evolution of genomics and underscore the importance of staying at the forefront of sequencing technologies for the accurate representation of complex genomes.

PacBio HiFi reads and Hi-C reads were used to generate a contig assembly with Hifiasm ^14^ producing a haplotype phased assembly. The contig level assembly produced single contigs for 9 chromosomes, while the remaining 3 chromosomes were each covered by 2 contigs each. Hi-C data were employed to hierarchically cluster the assembled contigs into 12 pseudo-chromosomes, by using the YaHS scaffolding tool ^15^. The T2T assembly had a single scaffold for each of the 12 pseudo-chromosomes and was larger in size than the corresponding IRGSP.10 genome (Figure 1a). The results of BUSCO analysis showed that the collapsed assembly covered 99.3% of universal single copy genes with an N50 of 30.7 Mb (Supplementary Table S1-3, Figure S2). For the two phased haplotypes (T2T); haplotype 1 covered 98.9% of the single copy orthologs with an N50 of 30.6 Mb, whilst haplotype 2 covered 94.9% single copy orthologs with an N50 of 29.3 Mb (Supplementary Table S4-9, Figure S3-4). The haplotype 1 chromosomes were larger than the haplotype 2 chromosomes (Figure1b). This first haplotype resolved Nipponbare genome, incorporated 3,050 new genes compared to IRGSP-1.0. and is expected to be a valuable and significant resource for rice researchers, and these additional genes had a wide range of functions (Supplementary Table S12). On these additional genes, 58 genes fell in new regions that were missing in the IRGSP genome, but most genes were in regions that were not new due to the improved accuracy of sequencing.

**Figure 1.**
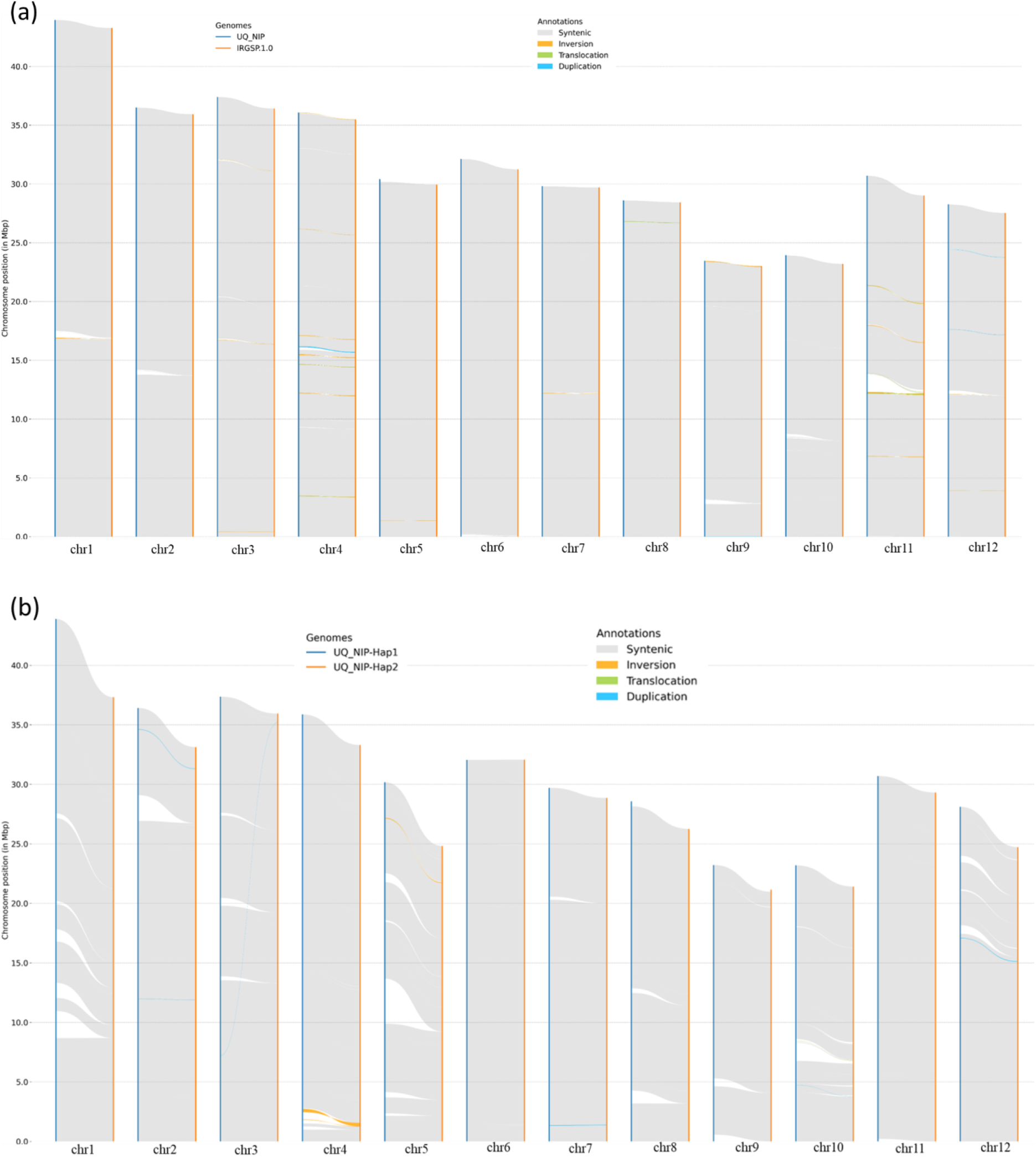
a) Sequence collinearity and structural variants (inversions, translocations, or duplications, and non-aligning regions) between UQ-NIP and IRGSP-1.0. The two assemblies were aligned using minimap2. Following alignment, the resulting BAM file was indexed using samtools and the detection of structural variations between these two genomes was carried out by using SyRI ^16-18^, b) The same approach was used to conduct a comparison of the genomic sequences between haplotype1 and hapotype2.

## Methods

### DNA extraction and sequencing

The CTAB method^19^ was used to extract DNA from young leaves of a rice (*Oryza sativa* cv Nipponbare) plant grown in a glasshouse at the University of Queensland. The high quality DNA extracted was sequenced using a PacBio (Pacific Biosciences) Sequel II platform to produce HiFi sequences.

### Genome assembly

Approximately 54.9 Gb of HiFi reads were obtained. HiC reads (59.6Gb) were downloaded from the NCBI Sequence Read Archive database (SRR6470741). De novo haplotype-resolved assembly of these reads was performed using hifiasm with parameters “--write-ec --write-paf -l0” ^14^. The contig assemblies were scaffolded using YaHS tool ^15^. QUAST and BUSCO were used to evaluate the quality and completeness of a genome assembly ^19,20^. A telomere identification toolkit (tidk) was used to search for tandem repeats of the telomeric sequence “TTAGGG” and “TAAACCC” and the exact location in the Nipponbare collapse, haplotype-1 and haplotype-2 assembly (https://github.com/tolkit/telomeric-identifier).

### Genome annotation

Repetitive DNA sequence were obtained from a Oryza repeat database ^21^and used to mask the genome with the Repeatmasker soft masking option ^22^. Protein sequences of viridiplantae from OrthoDB v.11 ^23^ and RNA-sequencing (RNA-seq) reads from the NCBI Sequence Read Archive database (SRR23560402, SRR23560417, SRR23560416, SRR23560419, SRR23560418, SRR23560409, SRR23107175, SRR23107177, SRR23107178, SRR8051554, SRR7974062, SRR8051550) were obtained. Quality and adapter trimmed RNA-seq reads were aligned to the masked genomes using HISAT2 ^24^. Annotation of protein-coding genes in Nipponbare was conducted using a combination of homology-based prediction, de novo prediction, and transcriptome-based prediction methods using Braker ^25^. BUSCO was used to assess the genome annotation completeness. The Large Gap Mapping tool (length fraction; 0.9 similarity fraction;0.9) of CLC was used to identify the new genes with the comparison of IRGSP-1.0 Nipponbare genes (CLC Genomics Workbench 23.0.05, QIAGEN, USA, http://www.clcbio.com) and further transcript evidence for these new genes was estimated (supplementary Table S11). The functional annotation of the identified additional genes was performed using OmicsBox 2.2.4 ^26^. CDS sequences were subjected to a BLASTX analysis with a specific e-value of 1.0E-10 against the non-redundant protein sequences database, utilizing Viridiplantae taxonomy. Subsequently, the CDS sequences were processed through InterProScan, and GO terms were extracted for all matches acquired via the BLAST search, employing Gene Ontology mapping with the Blast2GO annotation tool. The annotations generated from InterProScan and Blast2GO were then harmonized by merging the respective GO terms (Supplementary Table S12).

## Data availability

The whole genome sequence data reported in this paper have been deposited in the Genome Warehouse in National Genomics Data Center ^27,28^, Beijing Institute of Genomics, Chinese Academy of Sciences / China National Center for Bioinformation, under accession number GWHEQBP00000000 that is publicly accessible at https://ngdc.cncb.ac.cn/gwh.

## Acknowledgements

This research was supported by the ARC Centre of Excellence for Plant Success in Nature and Agriculture.

## Author Information

Authors and affiliations

### Contributions

M.A performed the experiments and analysed the data, A.F. and A.K. supervised and assisted in data analysis, P.O helped in writing the manuscript and data interpretation, R.H was involved in conception and supervision of the research. All authors approved the final version of the paper.

## Supplementary information

Supplementary Table S1-9 and Figures S1-4

