## Supplemental material Figures S1-4 Tables S1-10 for "An improved haplotype resolved genome reveals more rice genes"

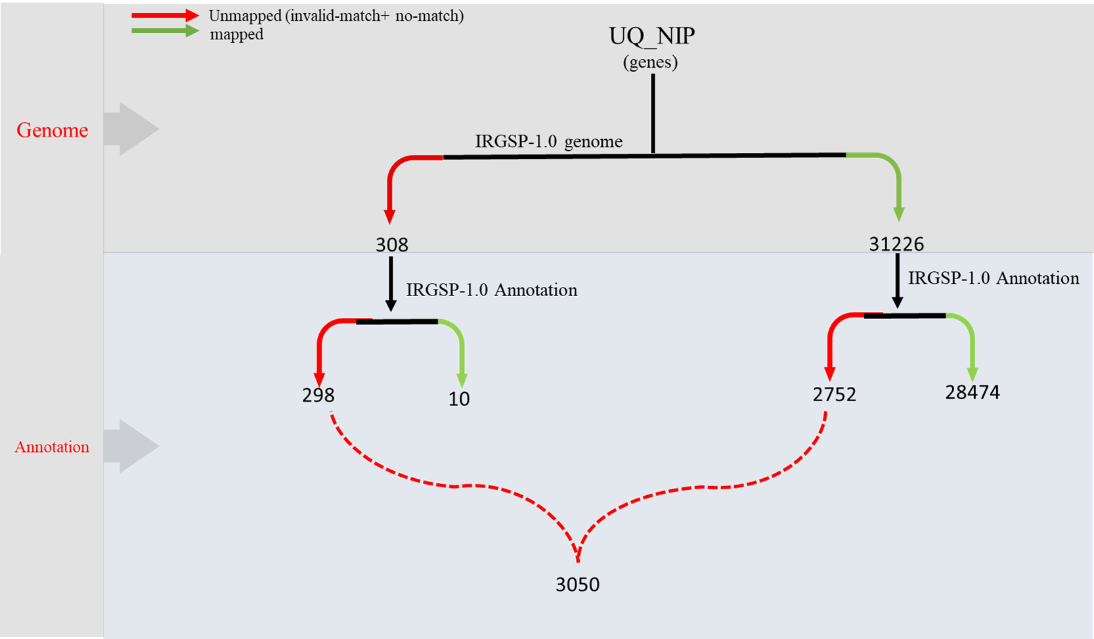

Figure S1. The Large Gap Mapping tool of CLC was used to identify the new genes by comparison of IRGSP-1.0 Nipponbare with UQ-Nipponbare. The genes from the UQ-Nipponbare were initially mapped to the IRGSP-1.0 Nipponbare genome sequence. After this initial mapping, both the mapped and unmapped genes were then subjected to another round of mapping, this time against the IRGSP-1.0 Nipponbare gene annotation.

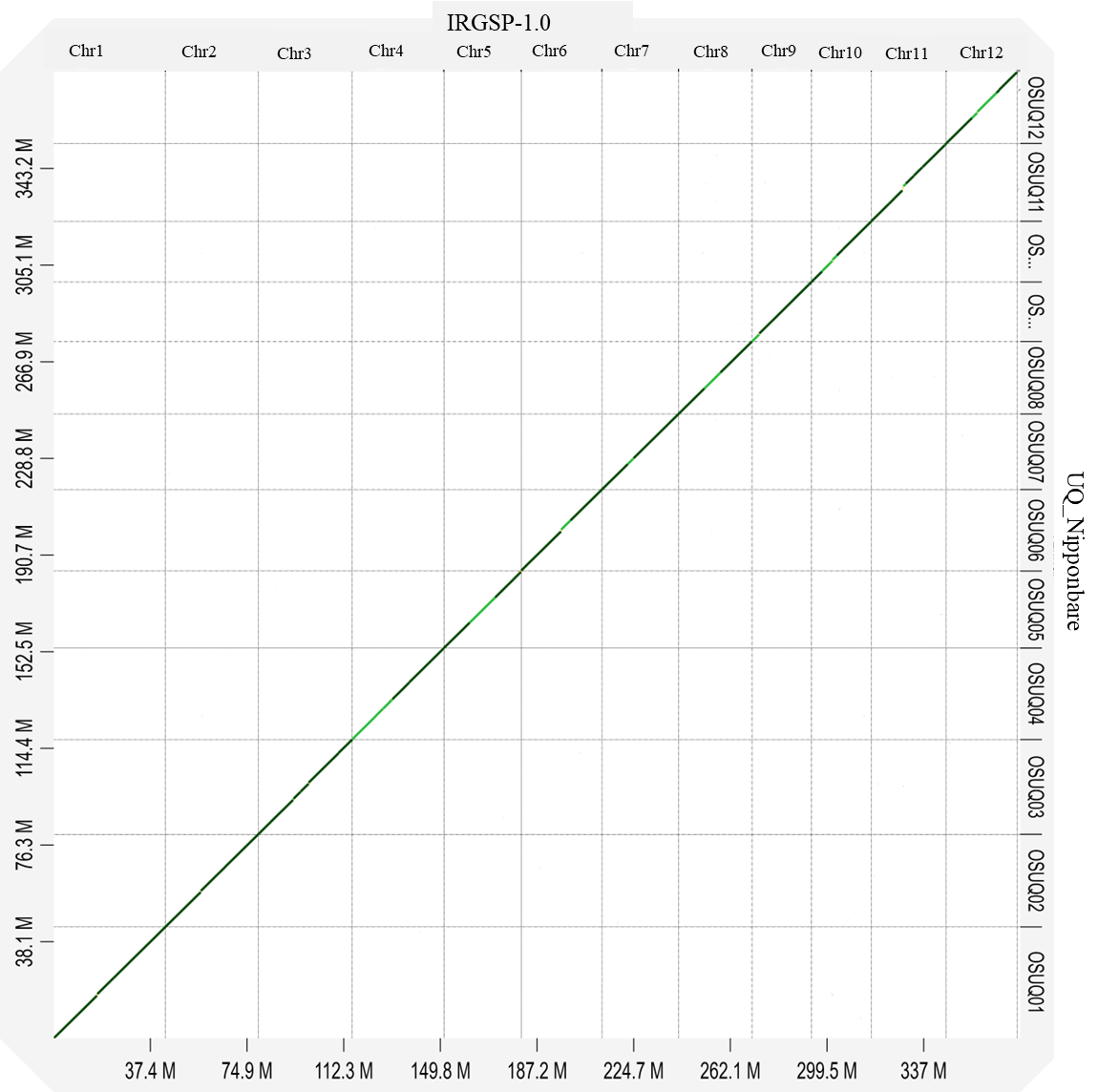

Figure S2. Dotplot of UQ_Nipponbare assembly against IRGSP.1.0 Nipponbare.

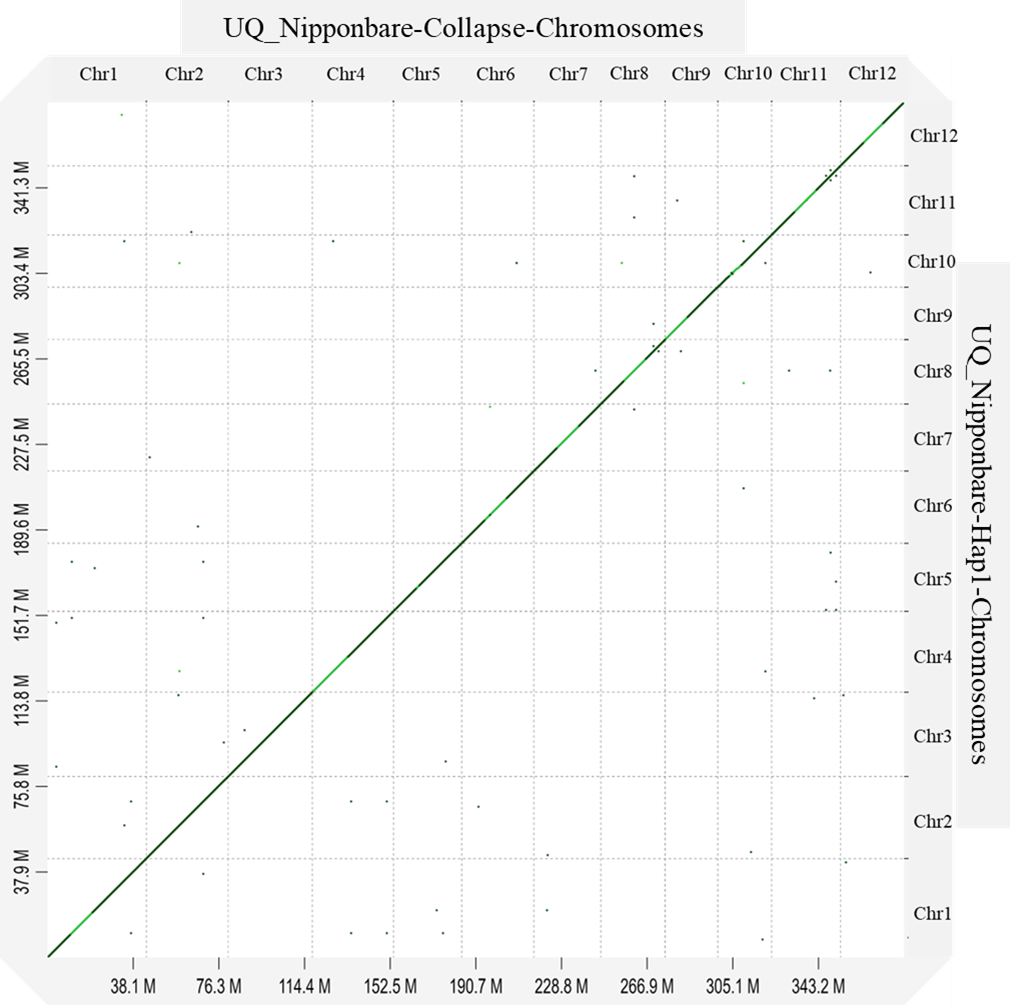

Figure S3. Dotplot of the chromosomes of the UQ_Nipponbare collapsed assembly against those of the UQ_Nipponbare haplotype 1 assembly.

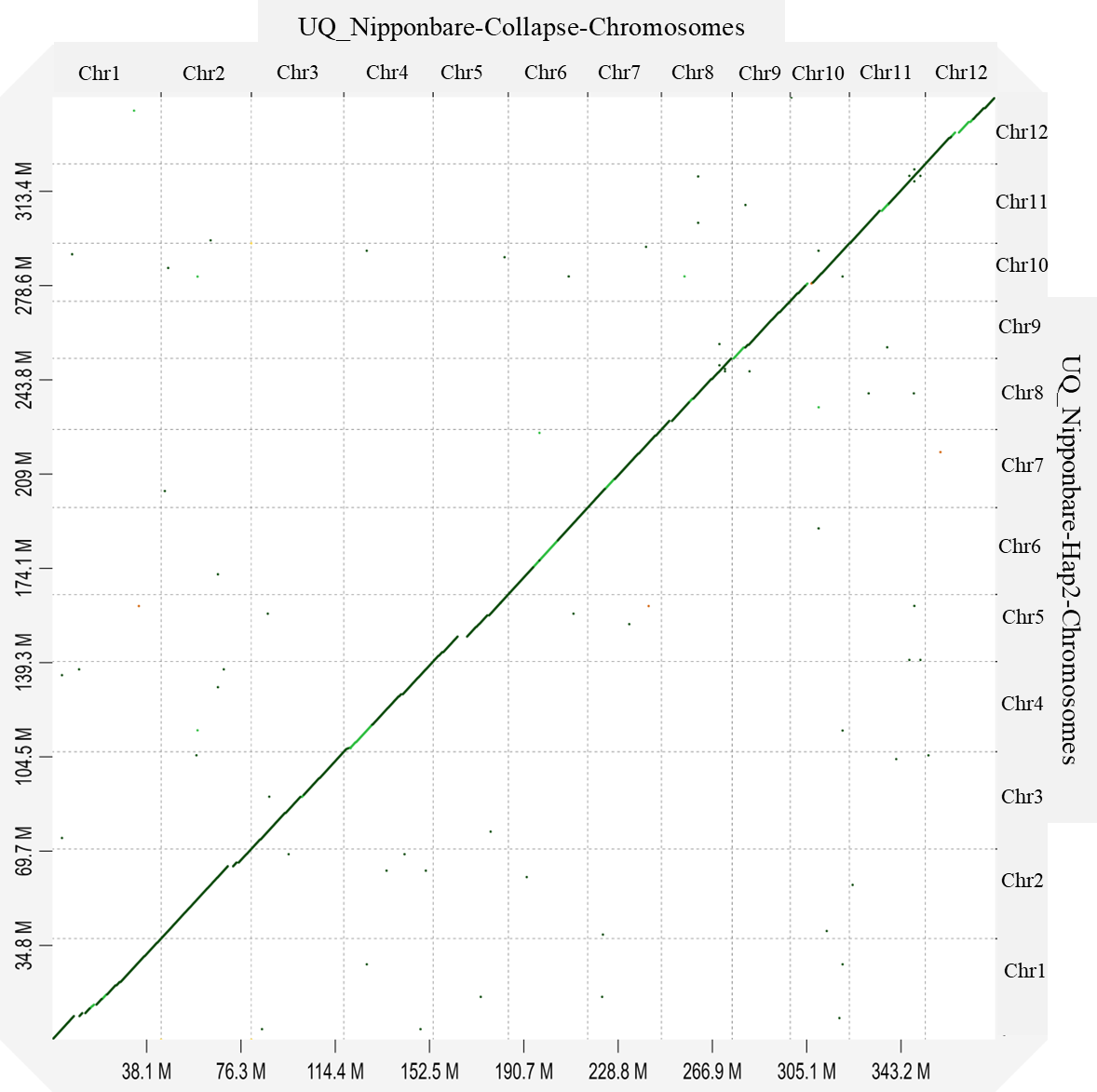

Figure S4. Dotplot of UQ_Nipponbare collapsed assembly chromosomes against UQ_Nipponbare haplotype 2 assembly.

Table S1. BUSCO statistics of the UQ-NIP-Collapsed Assembly.

| Statistics | Number | Percentage (%) |
| --- | --- | --- |
| Complete BUSCOs (c) | 422 | 99.3% |
| Complete and single-copy BUSCOs (S) | 412 | 96.9% |
| Complete and duplicated BUSCOs (D) | 10 | 2.4% |
| Fragmented BUSCOs (F) | 2 | 0.5% |
| Missing BUSCOs (M) | 1 | 0.2% |
| Total BUSCO groups searched | 425 | 425.00 |

Table S2. QUAST statistics of UQ-NIP-Collapsed Assembly

| # contigs (>= 0 bp) | 12 |
| --- | --- |
| # contigs (>= 1000 bp) | 12 |
| Total length (>= 0 bp) | 381,317,026 |
| Total length (>= 1000 bp) | 381,317,026 |
| # contigs | 12 |
| Largest contig | 43,960,277 |
| Total length | 381,317,026 |
| GC (%) | 44 |
| N50 | 30,712,252 |
| N90 | 23,948,751 |
| auN | 32,774,913 |
| L50 | 6 |
| L90 | 11 |

Table S3. UQ-NIP-Collapse d Assembly chromosome size and number of telomeres.

|  |  | Start-Telomere | | End-Telomere | |  |
| --- | --- | --- | --- | --- | --- | --- |
| #chr | length | seq | size | seq | size | Telomeres |
| OSUQ01 | 43,960,277 | TAAACCC | 8,471 | TTTAGGG | 12,360 | 2 |
| OSUQ02 | 36,506,049 | TAAACCC | 9,614 | TTTAGGG | 6,474 | 2 |
| OSUQ03 | 37,404,130 | TAAACCC | 1,718 | TTTAGGG | 9,043 | 2 |
| OSUQ04 | 36,083,220 | TAAACCC | 3,787 | TTTAGGG | 10,762 | 2 |
| OSUQ05 | 30,417,698 | TAAACCC | 6,265 | TTTAGGG | 7,541 | 2 |
| OSUQ06 | 32,132,640 | TAAACCC | 5,927 | TTTAGGG | 10,106 | 2 |
| OSUQ07 | 29,813,588 | TAAACCC | 10,228 | TTTAGGG | 7,120 | 2 |
| OSUQ08 | 28,607,546 | TAAACCC | 3,606 | TTTAGGG | 7,434 | 2 |
| OSUQ09 | 23,461,744 | TAAACCC | 5,710 |  |  | 1 |
| OSUQ10 | 23,948,751 | TAAACCC | 7,101 | TTTAGGG | 5,156 | 2 |
| OSUQ11 | 30,712,252 | TAAACCC | 246 | TTTAGGG | 4,299 | 2 |
| OSUQ12 | 28,269,131 | TAAACCC | 7,121 | TTTAGGG | 6,988 | 2 |

Table S4. BUSCO statistics of UQ-NIP-Hap1-Assembly.

| Statistics | Number | Percentage (%) |
| --- | --- | --- |
| Complete BUSCOs (c) | 420 | 98.9% |
| Complete and single-copy BUSCOs (S) | 410 | 96.5% |
| Complete and duplicated BUSCOs (D) | 10 | 2.4% |
| Fragmented BUSCOs (F) | 2 | 0.5% |
| Missing BUSCOs (M) | 3 | 0.6% |
| Total BUSCO groups searched | 425 | 425.00 |

Table S5. QUAST statistics of the UQ-NIP-haplotype1 Assembly.

| # contigs (>= 0 bp) | 12 |
| --- | --- |
| # contigs (>= 1000 bp) | 12 |
| Total length (>= 0 bp) | 379,234,557 |
| Total length (>= 1000 bp) | 379,234,557 |
| # contigs | 12 |
| Largest contig | 43,881,444 |
| Total length | 379,234,557 |
| GC (%) | 44 |
| N50 | 30,691,512 |
| N90 | 23,221,905 |
| auN | 32,642,297 |
| L50 | 6 |
| L90 | 11 |

Table S6. BUSCO statistics of UQ-NIP-haplotype 2 Assembly.

| Statistics | Number | Percentage (%) |
| --- | --- | --- |
| Complete BUSCOs (c) | 403 | 94.9% |
| Complete and single-copy BUSCOs (S) | 393 | 92.5% |
| Complete and duplicated BUSCOs (D) | 10 | 2.4% |
| Fragmented BUSCOs (F) | 2 | 0.5% |
| Missing BUSCOs (M) | 20 | 4.6% |
| Total BUSCO groups searched | 425 | 425.00 |

Table S7. QUAST statistics of the UQ-NIP-haplotype 1 Assembly.

| # contigs (>= 0 bp) | 12 |
| --- | --- |
| # contigs (>= 1000 bp) | 12 |
| Total length (>= 0 bp) | 348,265,595 |
| Total length (>= 1000 bp) | 348,265,595 |
| # contigs | 12 |
| Largest contig | 37,312,016 |
| Total length | 348,265,595 |
| GC (%) | 43 |
| N50 | 29,307,860 |
| N90 | 21,409,976 |
| auN | 29,954,189 |
| L50 | 6 |
| L90 | 11 |

Table S8. UQ-NIPhHaplotype 1 chromosome size and number of telomeres.

| Hap1-Chromosomes | Total size | Telomeres |
| --- | --- | --- |
| OSUQ01-hap1-01 | 43,881,444 | 2 |
| OSUQ02-hap1-02 | 36,408,562 | 2 |
| OSUQ03-hap1-03 | 37,357,616 | 2 |
| OSUQ04-hap1-04 | 35,866,358 | 2 |
| OSUQ05-hap1-05 | 30,178,946 | 2 |
| OSUQ06-hap1-06 | 32,049,832 | 2 |
| OSUQ07-hap1-07 | 29,708,413 | 2 |
| OSUQ08-hap1-08 | 28,566,876 | 2 |
| OSUQ09-hap1-09 | 23,221,905 |  |
| OSUQ10-hap1-10 | 23,192,984 | 2 |
| OSUQ11-hap1-11 | 30,691,512 | 2 |
| OSUQ12-hap1-12 | 28,110,109 | 2 |

Table S9. UQ-NIP-haplotype1 chromosome size and number of telomeres.

| Hap2-Chromosomes | Total size | Telomeres |
| --- | --- | --- |
| OSUQ01-hap2-01 | 37,312,016 | 2 |
| OSUQ02-hap2-02 | 33,128,268 | 2 |
| OSUQ03-hap2-03 | 35,934,962 | 2 |
| OSUQ04-hap2-04 | 33,298,206 | 2 |
| OSUQ05-hap2-05 | 24,819,415 | 2 |
| OSUQ06-hap2-06 | 32,067,105 | 2 |
| OSUQ07-hap2-07 | 28,849,953 | 2 |
| OSUQ08-hap2-08 | 26,254,248 | 1 |
| OSUQ09-hap2-09 | 21,159,188 | 1 |
| OSUQ10-hap2-10 | 21,409,976 | 2 |
| OSUQ11-hap2-11 | 29,307,860 | 1 |
| OSUQ12-hap2-12 | 24,724,398 | 2 |

Table S10. Summary of sequence data used for genome assembly.

| **Platform** | **PacBio (HiFi)** | |
| --- | --- | --- |
|  | SMRT cell 1 | SMRT cell 2 |
| Number of reads | 2.0M | 2.2M |
| Yield (bp) | 27.6 GB | 26.4 GB |
| Read quality (median) | Q33 | Q33 |
| Coverage | 72.40051191 | 69.18795232 |
